# Hematological and Neurological Expressed 1 (HN1): a predictive and pharmacodynamic biomarker of metformin response in endometrial cancers

**DOI:** 10.1101/602870

**Authors:** Nicholas W. Bateman, Pang-ning Teng, Erica Hope, Brian L. Hood, Julie Oliver, Wei Ao, Ming Zhou, Guisong Wang, Domenic Tommarello, Katlin Wilson, Kelly A. Conrads, Chad A. Hamilton, Kathleen M. Darcy, Yovanni Casablanca, G. Larry Maxwell, Victoria Bae-Jump, Thomas P. Conrads

## Abstract

Preoperative use of metformin in obese women with endometrioid endometrial cancer (EEC) reduces tumor proliferation and inhibits the mammalian target of rapamycin (mTOR) pathway, though is only effective in select cases. This study sought to identify a predictive and/or pharmacodynamic proteomic signature of metformin response to tailor its pharmacologic use. Matched pre- and post-metformin treated tumor tissues from a recently completed phase 0 window trial of metformin in EEC patients (ClinicalTrials.gov: NCT01911247), were analyzed by mass spectrometry (MS)-based proteomic and immunohistochemical analyses. Hematological and neurological expressed 1 (HN1) was significantly elevated in metformin responders (n=13) versus non-responders (n=7), which was also found to decrease in abundance in metformin responders following treatment; observations that were verified by immunohistochemical staining for HN1. Metformin response and loss of HN1 was assessed in RL95-2 and ACI-181 endometrial cancer cell lines. We further identified that silencing of HN1 abundance does not alter cellular response to metformin or basal cell proliferation, but that HN1 abundance does decrease in response to metformin treatment in RL95-2 and ACI-181 endometrial cancer cell lines. These data suggest that HN1 represents a predictive and pharmacodynamic biomarker of metformin response that, if validated in larger patient populations, may enable pre-operative EEC patient stratification to metformin treatment and the ability to monitor patient response.

**Novelty and Impact:** Hematological and Neurological Expressed 1 (HN1) is elevated in endometrioid endometrial cancer (EEC) patients that respond to metformin vs non-responders and decreases after treatment. HN1 decreases after metformin treatment of EEC cells in vitro, but is not necessary for metformin response or proliferation. We report HN1 as a novel predictive and pharmacodynamic biomarker of metformin response in EEC that may enable pre-operative EEC patient stratification to metformin treatment and the ability to monitor patient response.

## Introduction

Endometrial cancer (EC) is now the second most prevalent cancer among women in the USA and one of the few cancers for which disease incidence is on the rise, particularly for aggressive histologic subtypes (1) (2). Clinical management of EC includes total hysterectomy, bilateral salpingo-oopherectomy and pelvic and peri-aortic lymph node dissection and can be followed by adjuvant treatment with chemotherapy and vaginal brachytherapy. Given the increasing incidence and the paucity of effective treatments for advanced and recurrent EC, novel therapeutic agents that could be used alone or in combination with more traditional hormonal and cytotoxic chemotherapy to combat these trends are under active investigation. Emerging therapeutics for EC include the antihyperglycemic drug metformin (3-5). Owing to its effect of inhibiting hepatic gluconeogenesis, metformin is widely used clinically to manage type II diabetes. Metformin activates adenosine monophosphate kinase (AMPK) that, in turn, stimulates a number of catabolic pathways, such as glucose uptake, glycolysis and fatty acid oxidation and inhibits numerous ATP-consuming processes, such as fatty acid, cholesterol and protein synthesis (3,6). In the context of cancer, metformin has been shown to dramatically decrease proliferation of a number of different human cancer cell lines *in vitro* (5,7-9). Metformin has also been shown to effectively repress tumor growth in xenograft models of breast, prostate and colon cancer (10-12). In endometrial cancer cells, metformin-mediated AMPK activation decreases cell growth by inhibiting mTOR signaling (5).

Epidemiologic studies of diabetic patients have revealed those treated with metformin exhibit a significantly reduced risk to develop diverse cancers relative to non-metformin treated patients (13) (14). Focused meta-analyses have suggested that metformin improves disease outcome in diabetic cancer patients, as evidenced by a recent retrospective cohort study of 2,529 diabetic patients with early stage breast cancer that found that women receiving metformin and adjuvant chemotherapy had a higher complete pathologic response rate than those not taking metformin (15). In endometrial cancer, our team recently completed a preoperative phase 0 window study investigating the impact of short-term metformin treatment in women undergoing surgical staging for type I endometrial cancer who were obese (BMI >30) and/or have diabetes, but who were metformin treatment naïve (ClinicalTrials.gov: NCT01911247). We recently reported a comparative metabolomic analysis of serum and tumor tissues before and after metformin treatment stratified by response as measured by decreased abundance of the proliferation antigen Ki-67 in tumor tissues as measured by IHC staining (4). Metformin responders exhibited significantly decreased levels of phospho-AMPK as well as several downstream mTOR targets in tumor tissues and the serum metabolome reflected activated lipolysis suggesting activation of fatty acid metabolism following metformin treatment. We describe here a quantitative proteomics investigation of matched tumor tissues from the same cohort of pre- and post-metformin treated-patients collected during our phase 0 study of metformin in obese EC patients and identification of hematological and neurological expressed 1 (HN1) as a predictive and pharmacodynamic biomarker of metformin response in endometrial cancer patients.

## Methods

### Tissue Specimens

Formalin-fixed, paraffin embedded (FFPE) tissues were collected during pre-treatment biopsies or during primary surgery from endometrial cancer patients enrolled in a Phase 0 window trial for metformin treatment under IRB approved protocols (4). Thin (10 µm) tissue sections were cut using a microtome and placed on polyethylene naphthalate membrane slides. After staining with aqueous H&E, laser microdissection (LMD) was used to harvest tumor cells from the thin sections and were collected in 45 uL of LC-MS grade water (mean tumor cell area captured = ~16.1 ± 23 mm^2^).

### Liquid chromatography-tandem mass spectrometry (LC-MS/MS) proteomics

Tissue samples were processed and analyzed by high resolution LC-MS/MS on an Orbitrap Velos MS (Thermo Fisher, San Jose, CA) and HN1 IP samples (preparation described below) were similarly analyzed on a Fusion Lumos MS (Thermo Fisher) as previously described (16,17). Peptide identifications were filtered to include peptide spectrum matches (PSMs) passing an FDR < 1.0%. Proteins included in subsequent quantitative analyses were required to have a minimum of two PSMs. Significantly altered proteins were identified using edgeR (18) based on PSMs normalized to the patient sample exhibiting the lowest total PSM counts (edgeR p<0.05). Pathway analyses were performed using Ingenuity Pathway Analysis (Qiagen).

### Immunohistochemistry (IHC)

Rabbit polyclonal anti-HN1 antibody was purchased from Atlas Antibodies (Sigma-Aldrich - HPA059729, Bromma, Sweden). IHC was carried out in the Bond Autostainer (Leica Microsystems Inc. Norwell MA). Briefly, slides were dewaxed in Bond Dewax solution (AR9222) and hydrated in Bond Wash solution (AR9590). Antigen retrieval was performed for 20 min at 100ºC in Bond-Epitope Retrieval solution 1, pH-6.0 (AR9961). Slides were incubated with primary antibody (1:100) for 2 h. Antibody detection was performed using the Bond Polymer Refine detection system (DS9800). Stained slides were dehydrated and coverslipped. Positive and negative controls (no primary antibody) were included during the run.

### IHC image analysis

Stained slides were digitally scanned at 20× magnification using Aperio ScanScope-XT (Aperio Technologies, Vista, CA). Digital images were stored and analyzed within an Aperio eSlideManager Database. TMA images were segmented into individual cores using the Tissue Studio TMA portal (Tissue Studio version 2.7 with Tissue Studio Library v4.4.2; Definiens Inc., Munich, Germany). Epithelial cell enriched regions were digitally separated out for analysis using Tissue Studio Composer software (Definiens). Tissue Studio’s Nuclear Algorithm was then used to detect and enumerate cells that expressed HN1. The percentage of positive nuclei and an H-score (formula = (% at 1+) * 1 + (% at 2+) * 2 + (% at 3+) * 3) were determined for each TMA core.

### Cell culture and metformin treatment

RL95-2 was purchased from ATCC (Manassas, VA) and maintained in DMEM:F-12 medium supplemented with 10% fetal bovine serum, 1% penicillin-streptomycin and 0.005 mg/mL insulin (Sigma-Aldrich, St. Louis, MO). ACI-181 (19), a model of endometrioid endometrial cancer was a gift from Dr. John Risinger (Department of Obstetrics, Gynecology and Reproductive Biology, Michigan State University) and cultured in DMEM:F12 supplemented with 10% FBS 1% Pen/Strep. Both cell lines were maintained at 37 °C and 5% CO_2_. Metformin was purchased from Sigma-Aldrich.

### Cell Proliferation Assay

Cells were trypsinized with 0.25% trypsin-EDTA (ATCC), counted with 0.4% Trypan Blue using an automated cell counter (TC10, Bio-Rad, Hercules, CA) and viable cells were plated in 96-well plates at equivalent densities for each cell lines. For the metformin dose response assay, media was removed 24 hrs later and replaced with fresh media containing metformin (0, 1.56, 3.13, 6.25, 12.5, 25, 50, 100 mM) followed by incubation for 72h. Cell viability was assessed using the 3-(4,5-dimethylthiazol-2-yl)-5-(3-carboxymethoxyphenyl)-2-(4-sulfophenyl)-2H-tetrazolium (MTS) CellTiter 96 Aqueous One Solution Cell Proliferation Assay (Promega, Madison, WI) according to manufacturer’s instructions. Briefly, after a 3 h incubation at 37 °C, absorbance was measured at 490 nm using a microplate spectrophotometer (xMark, Bio-Rad). Two biological replicates were performed for each cell line with three technical replicates per metformin dose. For the cell proliferation assay, cells were plated in 96-well plates 72 h following transfection with siNT or siHN1 siRNA and cell viability was assessed daily by MTS assay as described above each day for four consecutive days for both the RL95-2 and ACI-181 cell lines.

### HN1 knockdown by siRNA

ON-TARGETplus Human JPT1 (HN1) siRNA SMART pool and ON-TARGETplus Human Non Targeting Control pool were purchased from Dharmacon. TransIT-siQUEST transfection reagent (Mirus Bio) was used for the siRNA transfections of the RL95-2 and ACI-181 cell lines. Cells were transfected with 75 nM siRNA according to the manufacturer’s protocol. Seventy-two hours post-transfection, cells were trypsinized, counted and seeded for cell viability assay. Cells were also harvested at 72 and 168 h after transfection to assess HN1 knockdown by immunoblotting and qPCR analyses. Two biological replicates were performed for each cell line.

### Quantitative PCR

cDNA was prepared from equivalent amounts of total RNA by reverse-transcription using the Superscript VILO cDNA synthesis kit (Invitrogen). HN1 (Hs00602957_m1) and GAPDH (Hs99999905_m1) TAQMAN assays were obtained from Applied Biosystems (Thermo Fisher). Quantitative PCR was performed using TAQMAN gene expression master mix of equivalent amounts of total cDNA for forty cycles (ABI GeneAmp 9700 DNA thermal cycler). Endpoint data was assembled by comparison of Delta-Ct values for HN1 versus corresponding GAPDH Delta-Ct values for each cell line. Triplicate technical replicates were performed.

### Immunoblot analyses

Cells were washed with cold PBS and lysed in lysis buffer containing 1% SDS and 150 mM NaCl (pH 7.4). Equivalent amounts of protein lysates were resolved via 4-15% mini-PROTEIN TGX gels (Bio-Rad) and transferred to PVDF membranes. Membranes were blocked for 1 hr using 5% non-fat dry milk (Bio-Rad) and incubated with primary antibody overnight at 4 °C, secondary antibody for 3 h at ambient temperature, and SuperSignal West Chemiluminescent Substrate (Thermo Scientific) for 5 min. Images were acquired using a ChemiDoc XRS+ system (Bio-Rad). Anti-Ki67 and anti-GAPDH were purchased from Abcam. Anti-MYC was purchased from Santa Cruz. Anti-HN1 was purchased from GeneTex. Anti-pAMPKα Thr172, anti-AMPKα, anti-pAKT1 Ser473, anti-AKT (pan) and goat anti-rabbit IgG HRP-linked were purchased from Cell Signaling Technologies.

### IP-MS analyses of the HN1

Twenty million RL95-2 cells were used per immunoprecipitation. For each immunoprecipitation, 30 µL of protein G sepharose 4 fast flow (GE Healthcare) was bound to 5 µL of anti-HN1 (GeneTex) for 2.5 h at 4°C with end-over-end rotation. Cells were lysed in 50 mM Tris (pH 7.4), 137 mM NaCL, 10% glycerol and 0.1% Triton X supplemented with 1 X Halt protease and phosphatase inhibitor (ThermoFisher Scientific) and 5mM EDTA for thirty min on ice. Lysate was spun at 16,000 × g for 20min at 4 °C and the supernatant was incubated with protein G sepharose 4 fast flow and anti-HN1 for 3 h at 4°C with end-over-end rotation. Sample was resolved on a 4-15% mini-PROTEAN TGX gel (Bio-Rad) and ten gel bands per lane were cut. Gel bands were digested using trypsin/LysC Mix (Promega) overnight and samples were cleaned up using Reversed-Phase ZipTip (Millipore) according to manufacturer’s protocol prior to LC-MS/MS analysis. Immunoprecipitation with anti-HN1 with protein G sepharose 4 fast flow was performed in duplicate. Immunoprecipitation with protein G sepharose 4 fast flow alone was used as a control.

### Public-Data Analysis

Normalized RNA-seq data (TCGA V2, (20) for HN1 and Ki-67 was downloaded from the https://gdac.broadinstritute.org for n=542 EC patients and transcript abundance was directly compared by Spearman correlation. Clinical characteristics were extracted from cgdsr (version 1.2.5) and a Kaplan-Meier analysis with log-rank testing was performed to assess the relationship between HN1 abundance and patient outcome using survival (version 2.37-7) package in R (version 3.1.2). For Kaplan-Meier curves, high versus low transcript expression was defined by the median cut-point capped at 60 months.

## Results

### Proteomic analysis of endometrial cancer (EEC) tumor tissues collected from pre and post-treated patient reveals conserved protein alterations between metformin responders and non-responders

Tumor tissues from endometrial cancer patients in a pre-operative phase 0 window trial were stratified as responders (n=13) or non-responders (n=7) to metformin treatment (Table 1). Response was defined as a decrease in IHC staining for Ki-67 when comparing pre-versus post-treatment endometrial cancer tissue as previously described (4). Quantitative LC-MS/MS-based global proteomic analyses of pathologically-defined, tumor cell populations harvested by laser microdissection from FFPE endometrial biopsies and endometrial cancer surgical tumor tissues identified 1,289 proteins by at least two peptide spectral matches (PSMs) across patients (Supplemental Table 1 and 2). Seventy-nine proteins were identified to be significantly altered (edgeR p-value ≤ 0.05) in pre-treatment tumor cells from metformin responder and non-responder patients (Figure 1 and Supplemental Table 3). Pathway analysis revealed top altered pathways to include adenosine monophosphate kinase (AMPK) signaling (Table 2, Supplemental Table 4A), along with those related to activating cellular signaling, regulating cellular proliferation, and inhibiting cell death and apoptosis in tissues in metformin responders (Table 2, Supplemental Table 4B). Protein alterations in pre-treatment tissues from responder versus non-responder patients (Supplemental Table 5A) were correlated with significant alterations in post versus pre-treated tissues from metformin responders (edgeR p-value ≤ 0.05), revealing 11 co-altered proteins between these groups (Uniprot Accession * and bolded, Supplemental Table 5A). Further analyses of metformin responders revealed activation of cell death and apoptosis signaling, but inhibition of viral infection as well as molecular transport in response to metformin (Supplemental Table 5B). Additional analyses of significant protein alterations in metformin non-responders (Supplemental Table 6A) also revealed activation of cell death as well as organ hypoplasia signaling, inhibition of T cell proliferation and leukocyte viability signaling in post versus pre-metformin treated tissues (Supplemental Table 6B).

**Table 1.** Metformin Window Trial Clinical Cohort (Adapted from Schuller KM et al, 2015).

| **Demographic** | **Responders**<br>**(n=13)** | **Non-Responders**<br>**(n=7)** | **P** |
| --- | --- | --- | --- |
| Age (years) | 60 (9.6) | 60 (9.6) | NS |
| BMI | 38.4 (5.4) | 41.8 (7.2) | NS |
| HgbA1c | 5.6 (0.4) | 5.6 (0.5) | NS |
| Duration of metformin<br>treatment (days) | 13.1 (5.3) | 17.4 (7.7) | NS |
| Stage |  |  |  |
| 1A | 8 | 7 | NS |
| 1B | 4 | 0 | NS |
| 2 | 1 | 0 | NS |
| Grade |  |  |  |
| 1 | 8 | 2 | NS |
| 2 | 4 | 3 | NS |
| 3 | 1 | 2 | NS |
| Underwent nodal<br>dissection | 11 | 7 | NS |

**Table 2.** Top canonical signaling pathways or disease and biofunctions activated or inhibited amongst 79 proteins significantly altered between metformin responders compared to non-responders.

| **Canonical Pathways** | **p-value** | **Proteins (#)** |
| --- | --- | --- |
| Glucocorticoid Receptor Signaling | 2.75E-05 | 8 |
| BER pathway | 8.13E-04 | 2 |
| AMPK Signaling | 1.05E-03 | 5 |
| Mismatch Repair in Eukaryotes | 1.45E-03 | 2 |
| Aryl Hydrocarbon Receptor Signaling | 1.62E-03 | 4 |

| **Functional Annotation** | **p-value** | **Activation z-score** | **Proteins (#)** |
| --- | --- | --- | --- |
| Viral Infection | 1.18E-03 | 2.776 | 18 |
| Cell proliferation of tumor cell lines | 1.00E-06 | 2.702 | 25 |
| Repair of DNA | 1.04E-04 | 2.599 | 8 |
| Migration of cells | 7.75E-06 | 2.265 | 26 |
| Proliferation of prostate cancer cell lines | 1.63E-04 | 2.157 | 8 |
| Death of embryo | 2.87E-06 | -1.937 | 7 |
| Neural tube defect | 3.74E-03 | -1.949 | 4 |
| Apoptosis | 2.15E-07 | -2.164 | 34 |
| Cell death of central nervous system cells | 2.39E-03 | -2.173 | 6 |
| Cell death | 1.28E-10 | -2.288 | 45 |

**Figure 1.**
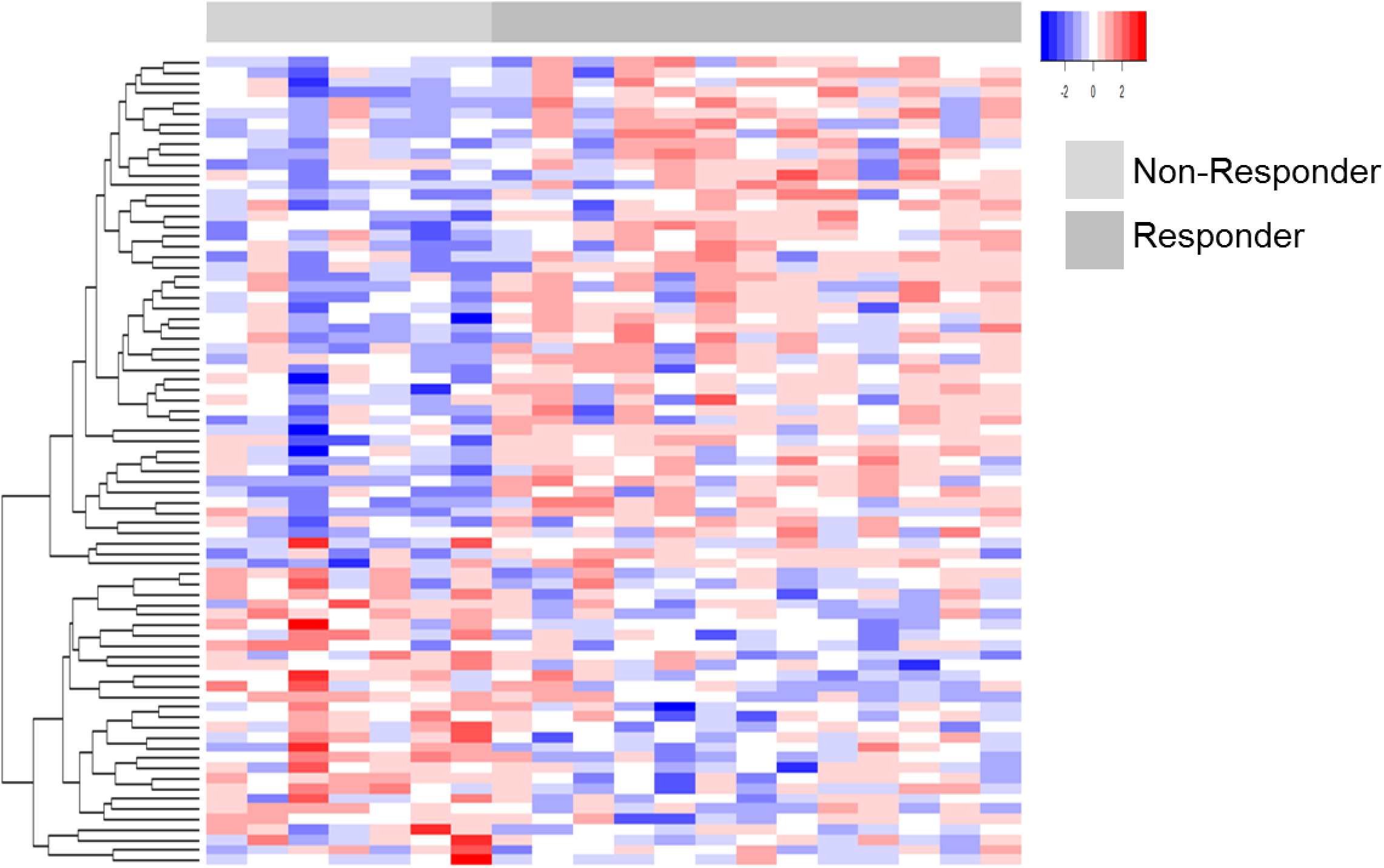
Differential proteomic analyses of endometrial cancer (EC) tissues collected from women who responded (n=13) or did not respond (n=7) to metformin treatment. Heatmap details a supervised analyses of 79 proteins (edgeR p<0.05) significantly altered between EC patients who did or did not respond to metformin treatment.

### Hematological and Neurological Expressed 1 (HN1) - a predictive and pharmacodynamic biomarker of metformin response in endometrial cancers

Among proteins correlating with response, we prioritized Hematological and Neurological Expressed 1 (HN1) for further validation as this protein exhibited the greatest fold-change difference in pre-metformin treatment biopsies from responders versus non-responders (log_2_ ratio = 3.84, edgeR p-value = 9.78E-6), and was decreased in abundance in post versus pre-treated tumor tissues in metformin responders (log_2_ ratio = −1.06, edgeR p-value = 0.02), but remaining largely unaltered in non-responders (Figure 2). Immunohistochemical staining for HN1 in the same set of patient tissues verified the proteomics data (Figure 3, Supplemental Table 7). These results confirmed that HN1 expression is significantly elevated in responder versus non-responder tissues, pre-metformin treatment (p-value = 0.05) and that HN1 expression was decreased in responders, but unaltered in non-responders, post-metformin treatment (p-value = 0.011) (Figure 3B).

**Figure 2.**
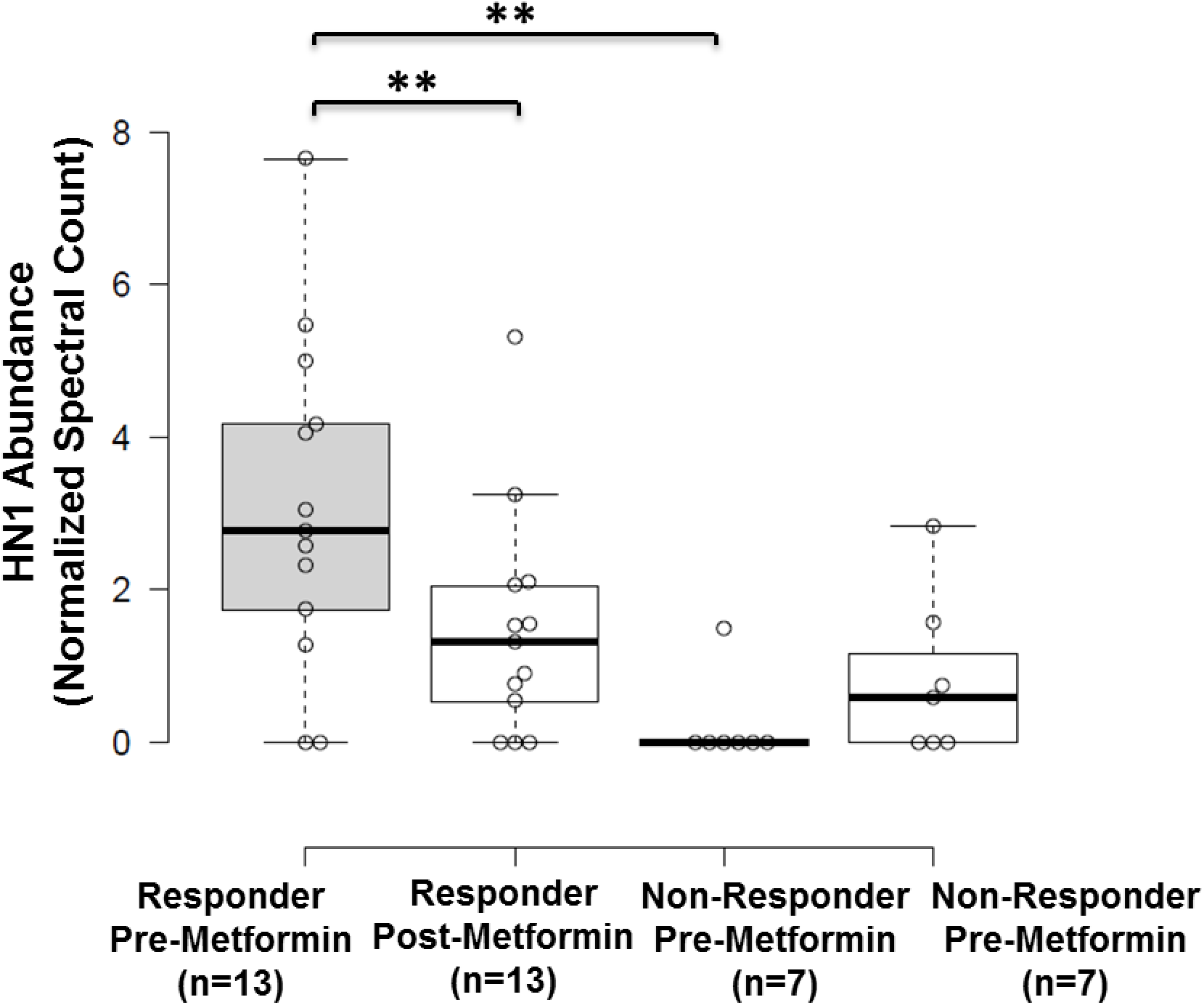
Hematological and neurological expressed 1 (HN1) protein in elevated in metformin responders and decreases following metformin treatment. HN1 was observed as significantly abundant in metformin responder versus non-responder patients and was significantly decreased in abundance following metformin treatment in responders by LC-MS/MS-based proteomic analyses (** edgeR p<0.05).

**Figure 3.**
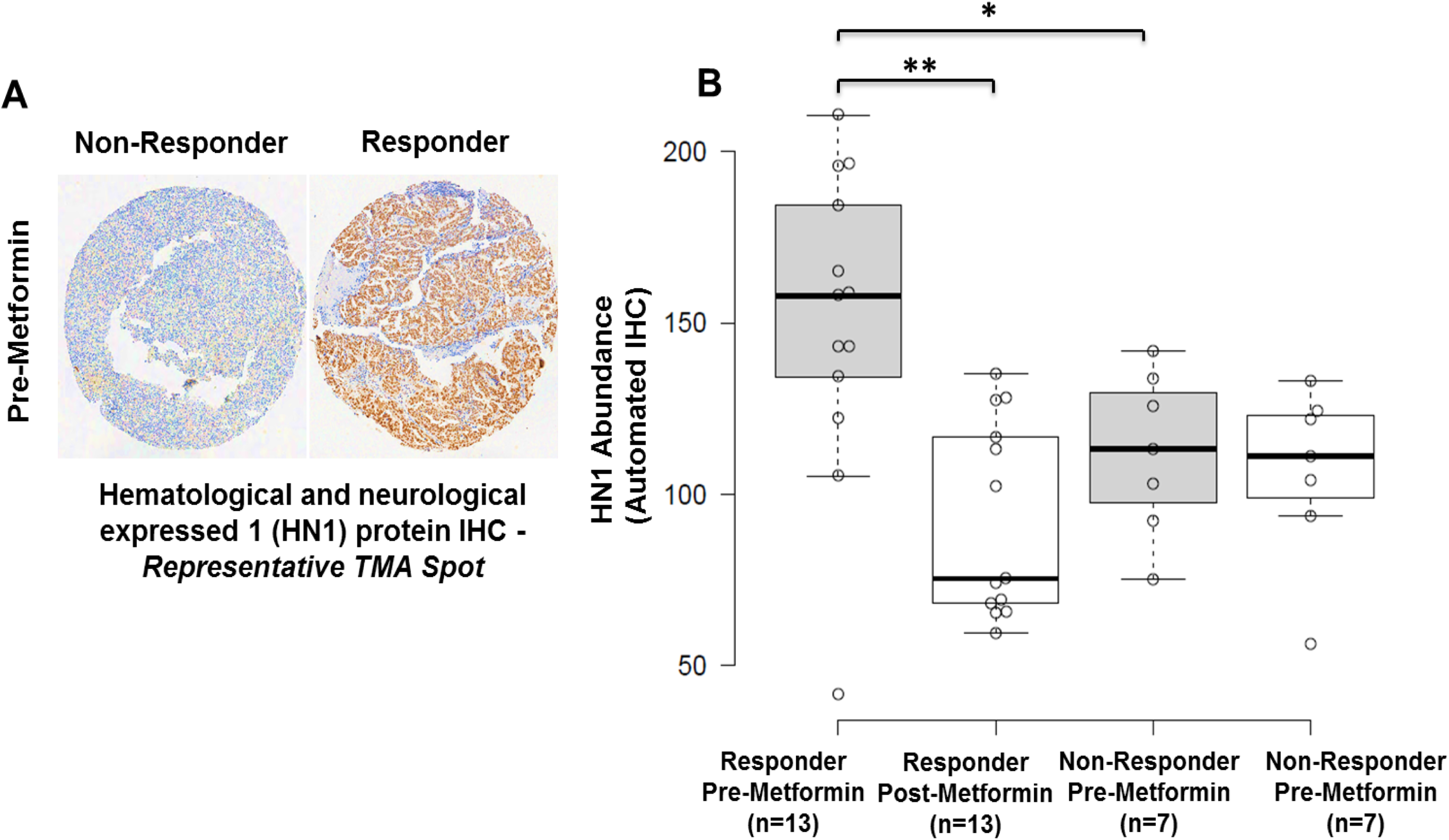
Verification of altered HN1 protein abundance by immunohistochemical analyses of EEC patients tissues collected from metformin responder (n=13) versus non-responders (n=7). A: HN1 IHC staining in EEC tissues harvested from a representative responder versus non-responder patient, pre-metformin treatment. B: IHC H Score for HN1 abundance in metformin responders and non-responder, pre and post-Metformin treatment. * p-value = 0.05, ** p=value = 0.011.

### Hematological and Neurological Expressed 1 (HN1) expression is decreased following metformin treatment of endometrial cancer cells, but HN1 is not necessary for response to metformin or for endometrial cancer cell proliferation

The endometrial cancer cell lines RL95-2 and ACI-181 were treated with 20 mM metformin (~LD50) for 96 and 120 h and HN1, AMPKα, p-AMPKα (T172), and Ki-67 protein abundance was assessed by immunoblotting. Metformin induced activation of AMPKα, as evidenced by increase in p-AMPKα (T172) abundance and further mediated a decrease in Ki-67 and HN1 abundance (Figure 4). We further assessed the impact of HN1 on response to metformin in EC cells in which HN1 expression was silenced by small interfering RNAs targeting HN1 mRNA (Figure 5A, 5B, 5D and 5E). These analyses revealed that loss of HN1 expression did not alter the response of EC cells to metformin treatment (Figure 5C and 5F). As AKT has previously been noted to be hyperactivated in low HN1 backgrounds (21), we further assessed activation of AKT in HN1 silenced cells. However, we did not observe alterations of p-AKT (S473) abundance in EC cells transfected with HN1-specific versus non-targeting siRNAs (Figure 5A and 5D). Further, recent evidence has shown that HN1 knockdown results in decreased abundance of the MYC oncogene (22). We further assessed MYC abundance in HN1 silenced endometrial cancer cells, but did not reproduce this finding (Figure 5A and 5D). We further assessed the impact of silencing HN1 on the proliferation rate of RL95-2 and ACI-181 cells (Figure 6). These analyses revealed that loss of HN1 expression does not significantly alter the proliferation of endometrial cancer cells.

**Figure 4.**
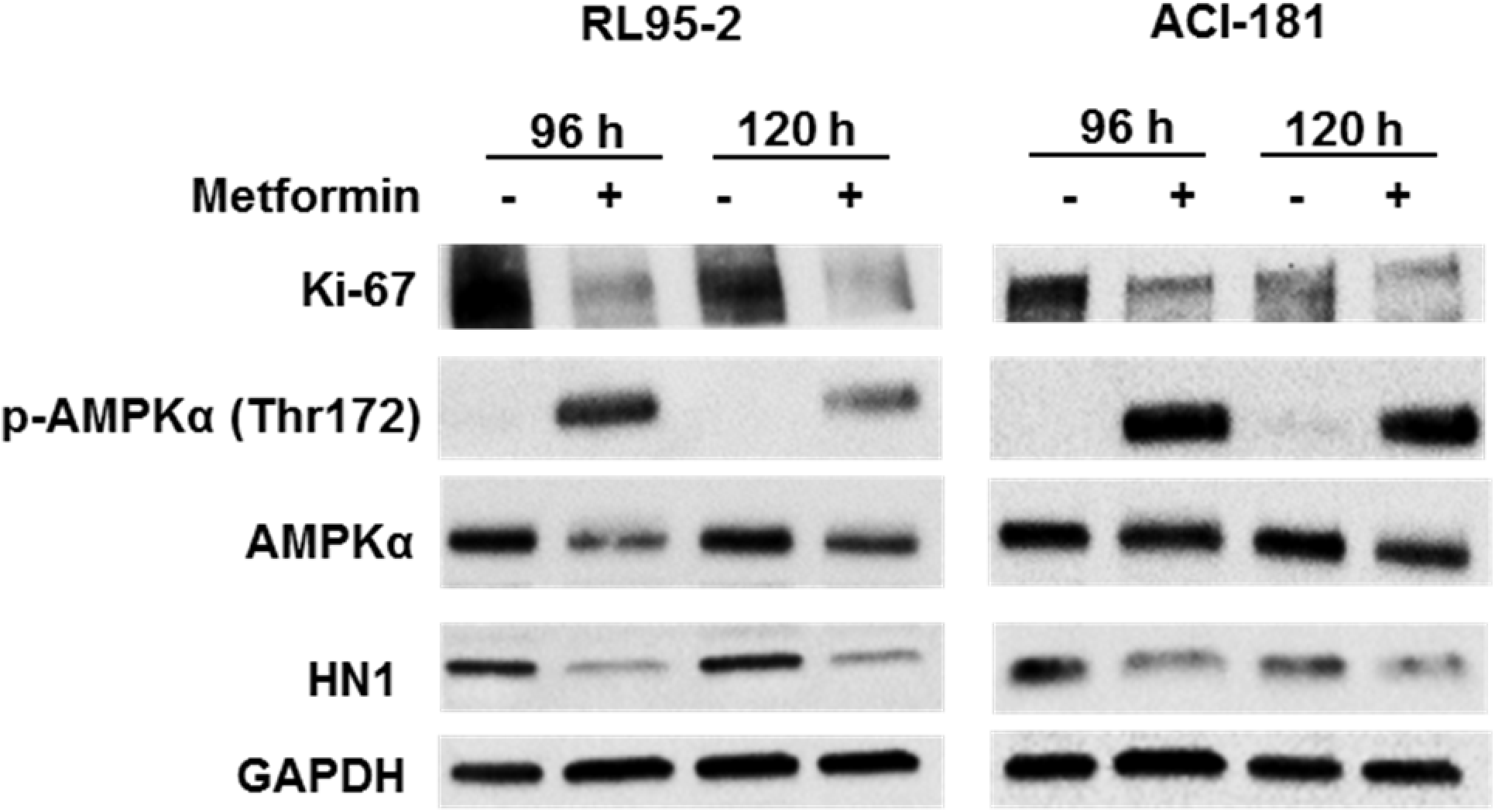
HN1 decreases in response to metformin treatment in endometrial cancer cells. RL95-2 and ACI-181 cells were treated with metformin (20mM) for 96h or 120h and equivalent amounts of protein lysate was immunoblotted for HN1 protein abundance as well as AMPKα, p-AMPKα (Thr172), and Ki-67 proteins.

**Figure 5.**
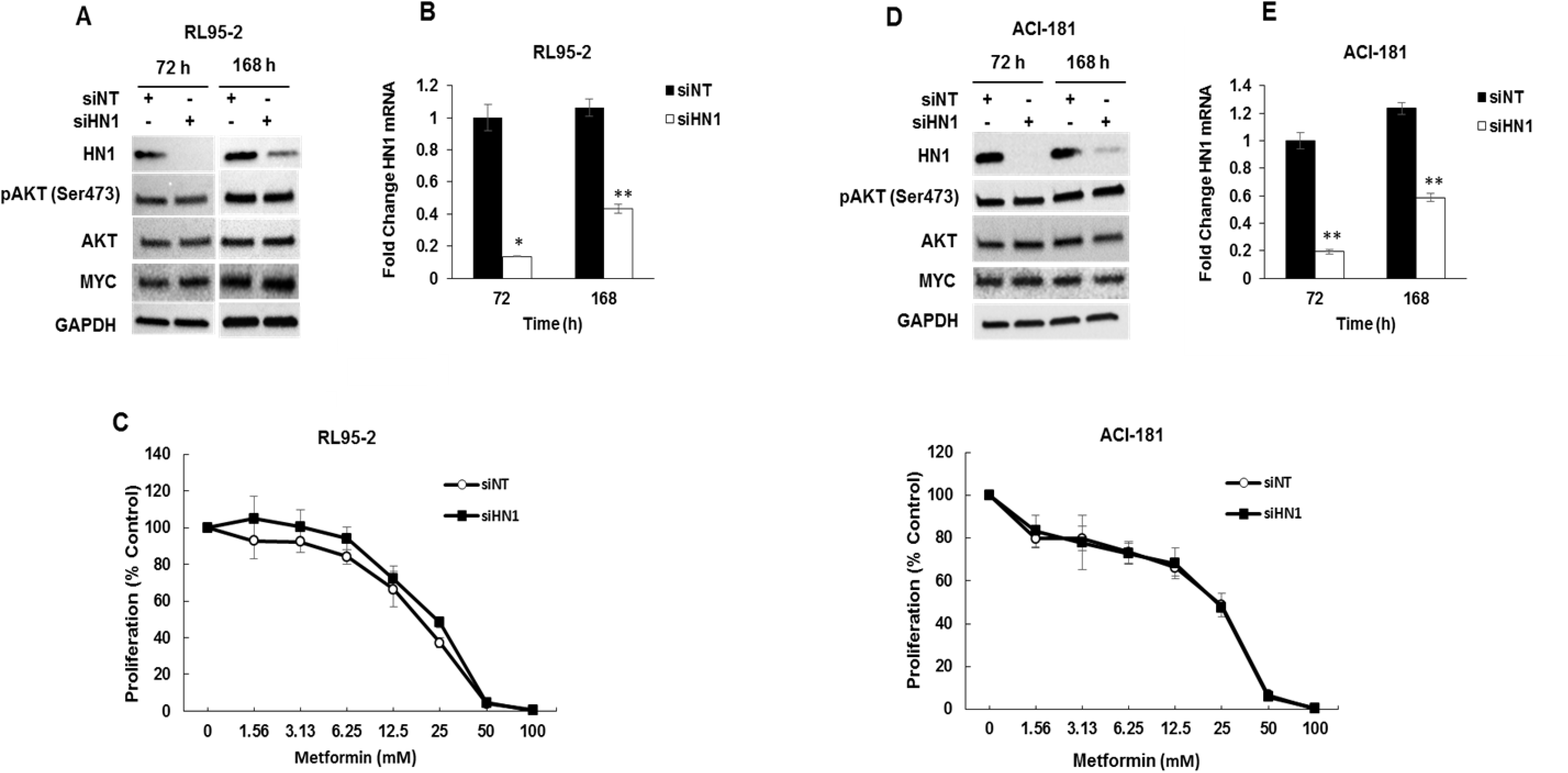
HN1 knockdown does not alter response of endometrial cancer cell lines to metformin. HN1 expression was knocked down in RL95-2 and ACI-181 cells by small interfering RNA (siRNA) targeting HN1 or with a non-targeting (siNT) control and confirmed at the protein (A & D) and mRNA (B & E) levels before (72h) and following (168h) dose-response analyses with metformin treatment; p-value was determined by student t-test. **p < 0.001, *p < 0.005. RL95-2 and ACI-181 cells were treated with metformin 72h following siRNA transfection and dose-response was assessed after an additional 72h by MTS assay.

**Figure 6.**
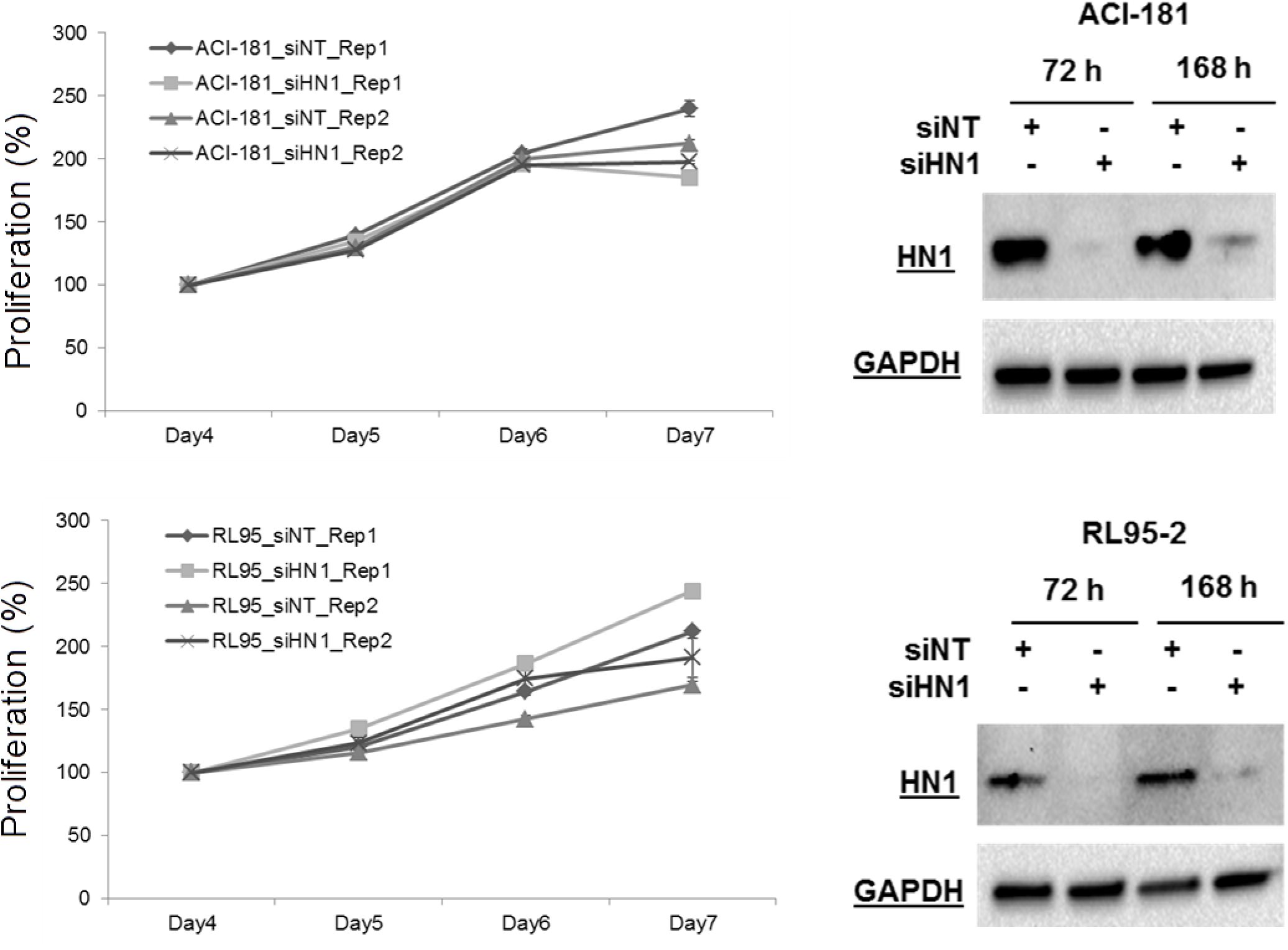
HN1 knockdown does not alter the proliferation of endometrial cancer cell lines. HN1 expression was silenced in RL95-2 and ACI-181 cells by small interfering RNA (siRNA) targeting HN1 or with a non-targeting (siNT) control and confirmed at the protein level (B & D) before (72h) and following (168h) daily assessment of cellular proliferation by MTS assay; p-value was determined by student t-test. Each data point reflects triplicate technical replicates measured at two sub-confluent cell densities per day, error bars reflect standard error. Rep 1 = biological replicate #1, Rep2 = biological replicate 2.

### Hematological and Neurological Expressed 1 (HN1) is correlated with Ki-67 expression and associated with altered overall survival in endometrial cancer patients

As Ki-67 and HN1 abundance were both decreased following metformin treatment, we assessed correlation trends of HN1 protein abundance (Supplemental Table 7) relative to Ki-67 protein abundance previously assessed in these tissues (4). We found that HN1 and Ki-67 exhibit concordant abundance trends in comparisons of both pre-(Spearman = 0.44) as well as in post-(Spearman=0.58) metformin treated tissues. We further assessed correlation of HN1 and Ki-67 transcript expression using a public RNA-seq dataset established from n=542 endometrial cancer patient tumor tissues (20) and found that these genes were positively correlated in endometrial cancer patient tissues (Spearman = 0.37, Supplemental Figure 1A). We also assessed whether HN1 abundance was directly correlated with disease outcome in endometrial cancer patients and found that elevated expression of HN1 was associated with poor overall survival (n=540 EC patients, Log-rank p-value = 0.047, Supplemental Figure 1B) but not with disease-free survival (n=499 EC patients, Log-Rank p-value = 0.14, data not shown).

### Hematological and Neurological Expressed 1 (HN1) exhibits binding partners regulating metabolic signaling pathways in endometrial cancer cells

We performed an immunoprecipitation and mass spectrometry (IP-MS)-based analyses of HN1 in subconfluent RL95-2 cells to identify putative binding partners of HN1 (Supplemental Table 8A). HN1 was the most abundantly identified protein in HN1 IP-MS biological replicates analyses, i.e. 61 ± 8 PSMs, following removal of proteins identified in a companion control IP-MS analyses of sepharose beads alone that contained no anti-HN1 antibody. Top functional pathways enriched following analyses of proteins abundantly identified with HN1, i.e. ≥ 10 PSMs in both HN1 IP-MS analyses (designated with * in Supplemental Table 8A) were associated with metabolic pathways such as purine nucleotides de novo biosynthesis, glycolysis and gluconeogenesis as well as fatty acid biosynthesis signaling (Supplemental Table 8B).

## Conclusions

The current study sought to identify proteomic alterations in tumor tissues harvested from endometrial cancer patients before and after daily treatment with metformin (850mg) for 4 weeks prior to surgical staging. Women who responded to metformin exhibited a significant decrease (7 to 50%) in expression of the cellular proliferation marker Ki-67 in post-treated tumor tissues, whereas non-responders exhibited a slight increase in Ki-67 IHC staining (~2-12%), as described previously (4). A global quantitative LC-MS/MS-based proteomics analysis identified discrete protein alterations in tumor cells harvested by LMD from pre-treated endometrial biopsies that correlate with canonical cellular pathways that include AMPK signaling, a kinase regulating cellular energy homeostasis activated by metformin (6), as well as pathways involved in activating cellular proliferation and inhibiting cell death signaling in metformin responders. Among the proteins altered, Hematological and Neurological Expressed 1 (HN1) was the most significantly elevated protein in pre-treated endometrial biopsies from metformin responders (responder versus non-responder logFC=3.42, edgeR p-value = 9.78E-06). We cross-correlated HN1 in pre and post-treatment biopsies in metformin responders and found that HN1 was also significantly decreased following metformin treatment (post-versus pre-treatment logFC = −1.06, edgeR p-value = 0.02) – a finding that was independently verified by IHC.

Hematological and Neurological Expressed 1 (HN1), also known as Jupiter microtubule associated homolog 1 (JPT1), is a ~16 kDa protein (Q9UK76, uniprot.org) initially identified as an abundant transcript in both murine hematopoietic and brain cells (23). As HN1 was significantly elevated in patients who responded to metformin, we assessed the impact of silencing HN1 expression on response to metformin using two endometrial cancer cell line models, i.e. RL95-2 and ACI-181 (Figure 5). We found that loss of HN1 did not alter cellular response to metformin treatment *in vitro*. In cancer, HN1 has been shown to inhibit the growth of androgen-sensitive, but promote the growth of androgen-insensitive, prostate cancer cells through altered androgen receptor stability and by modulating cell metabolism and cell cycle progression through protein kinase B (AKT) and downstream glycogen synthase kinase-3 beta (GSK3β)/ β-catenin (CTNNB1)-dependent mechanisms (21,24). Specifically, silencing of HN1 expression increased activation of AKT via upregulation of p-AKT (S473) in androgen-dependent and independent prostate cancer cell lines (21). We similarly assessed modulation of p-AKT (S473) following knockdown of HN1 in endometrial cancer cell lines, though we did not reproduce this finding (Figure 4). Recent evidence has also shown that elevated HN1 expression is directly correlated with poor prognosis in breast cancer patients (22). Further investigations revealed that HN1 increases the migratory and invasive potential of breast cancer cells by upregulating the MYC oncogene and downstream effectors *in vitro* and further increases breast tumorigenesis *in vivo* (22). We, however did not see any alterations in MYC protein abundance following silencing of HN1 in endometrial cancer cells (Figure 5). We found that metformin treatment inhibited endometrial cancer cell proliferation *in vitro*, as evidence by decreased Ki-67 levels, and that this response was further accompanied by loss of HN1 abundance (Figure 5). Pathway analyses of significantly altered proteins in metformin pre-treatment tumors from responders revealed that cellular proliferation signaling was altered in metformin responders as evidenced by elevation of proliferating cell nuclear antigen in responder patients (PCNA, edgeR LogFC=1.29, p-value = 0.025). PCNA regulates DNA replication via interactions with DNA polymerase delta and is a marker of cellular proliferation often assessed during routine histopathology analyses of tumor tissues along with Ki-67 to determine the proliferative indices of tumor cells *in situ* (25). Indeed, we previously observed (4) that pre-treatment tumor tissues from metformin responders exhibited elevated Ki-67 expression levels versus non-responders (47.3% vs. 24.9%, p-value = 0.004), providing suggestive evidence that these patients may harbor tumor cells with a greater proliferative capacity. That we do not observe an association between HN1 and response to metformin and as HN1 levels closely parallel PCNA levels in responders versus non-responders, HN1 abundance may also serve as a surrogate measure of proliferative activity in endometrial cancer cells. Similarly to our findings with HN1, recent assessments of Ki-67 have identified that this hallmark biomarker of cellular proliferation does not directly contribute to tumor cell proliferation (26,27), but rather functions to maintain cancer stem cell populations (27) and facilitates chromosomal motility and mitotic spindle interactions (28). To gain further insight into the possible biologic roles of HN1 in endometrial cancer cells, we performed an immunoprecipitation-mass spectrometry based analyses of HN1 in sub-confluent RL95-2 cells to identify functional binding partner candidates. These analyses revealed putative HN1 binding proteins to be predominantly associated with metabolic signaling pathways such as nucleotide and aerobic glycolysis and fatty acid signaling. Future studies will be focused on validating these HN1 binding partner candidates and elucidating the roles of these targets along with HN1 in the regulation of metabolic signaling pathways as they pertain to metformin treatment.

Immunohistochemical markers of metformin response used in clinical trials to date include assessments of AMPK abundance and activation state (p-AMPK, T172), mTOR signaling, such as p-S6 kinase as well as p-EIF4-BP1, as well as assessing markers of cellular proliferation (29). Indeed, we observe activation of AMPK via increased levels of p-AMPK (T172) in endometrial cancer cells treated with metformin *in vitro* (Figure 4). We did not, however, observe elevated p-AMPK (T172) by IHC in endometrial cancer tissues following metformin treatment (4), though Ki-67 was clearly altered these same tissues, recapitulating our findings *in vitro*. Comparative analyses of Ki-67 and HN1 protein abundance by IHC analyses in pre and post-metformin treated patient tissues revealed these protein abundances to be positively correlated. We assessed the correlation of HN1 and Ki-67 mRNA abundance in public data for over 500 endometrial cancer patients and found that HN1 and Ki-67 expression are positively correlated (R=~0.365). Furthermore, we also identified that elevated HN1 is associated with poor overall survival, underscoring that high HN1 expressing patients may experience a survival benefit from metformin treatment. With further validation, these data suggest that IHC assessment of HN1 in pre-surgical endometrial tissue biopsies may aid to prioritize patients for neoadjuvant treatment with metformin, and that this protein may further function as a pharmacodynamic surrogate of therapeutic response. Future studies will focus on validating the performance of HN1 as a pharmacodynamic biomarker of metformin response in an expanded cohort of endometrial cancer patients that will include both obese and non-obese patients.

## Abbreviations

HN1: Hematological and Neurological Expressed 1
JPT1: Jupiter microtubule associated homolog 1
EC: endometrial cancer
EEC: cndometrioid endometrial cancer
Ki-67: Antigen Ki-67
MS: mass spectrometry
MS/MS: tandem mass spectrometry
LC: liquid chromatography
mTOR: mammalian target of rapamycin
AMPK: adenosine monophosphate kinase
ATP: adenosine triphosphate
LMD: laser microdissection
H&E: hematoxylin and eosin
IHC: Immunohistochemistry
PSMs: peptide spectral matches
FDR: false discovery rate
TMA: tissue microarray
CO2: carbon dioxide
siRNA: small interfering RNA
mM: millimolar
MTS: 3-(4,5-dimethylthiazol-2-yl)-5-(3-carboxymethoxyphenyl)-2-(4-sulfophenyl)-2H-tetrazolium
siNT: small interfering non-targeting RNA
siHN1: small interfering HN1 RNA
qPCR: quantitative polymerase chain reaction
cDNA: complimentary DNA
DNA: deoxyribonucleic acid
RNA: ribonucleic acid
GAPDH: glyceraldehyde 3-phosphate dehydrogenase
SDS: sodium dodecyl sulfate
NaCl: sodium chloride
PVDF: polyvinylidene difluoride
AKT: protein kinase B
IgG: immunoglobulin
HRP: horseradish peroxidase
IP-MS: immunoprecipitation mass spectrometry
EDTA: ethylenediaminetetraacetic acid
TCGA: The Cancer Genome Atlas
RNA-seq: RNA sequencing
LD50: lethal dose 50
mRNA: messenger RNA
MYC: MYC oncogene
Mg: milligram
logFC: Log fold-change
CTNNB1: beta catenin
S473: serine 473
PCNA: proliferating cell nuclear antigen
p-S6 kinase: phosphorylated ribosomal protein S6 kinase beta-1
p-EIF4-BP1: phosphorylated Eukaryotic translation initiation factor 4E-binding protein 1

## Disclaimer

The views expressed herein are those of the authors and do not reflect the official policy of the Department of Army/Navy/Air Force, Department of Defense, or U.S. Government.

---

**Supplemental Figure 1.**
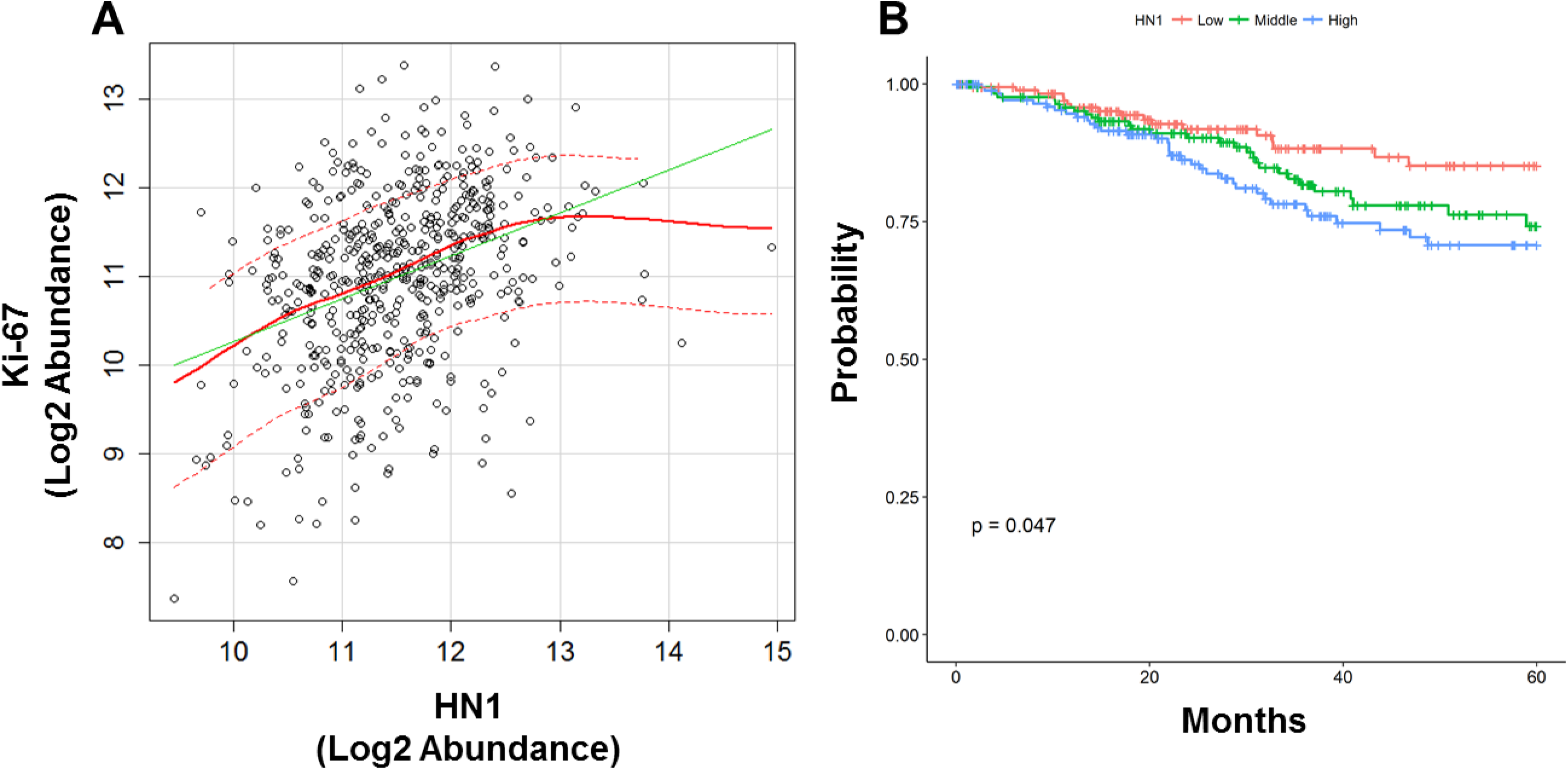
HN1 gene expression correlates with Ki-67 expression and was associated with altered overall survival in endometrial cancer patients. A: Correlative HN1 and Ki-67 mRNA abundance determined by RNA-seq analyses of tumor tissues from endometrial cancer patients (n=542, The Cancer Genome Atlas); Spearman = 0.37, Pearson = 0.36. B: Overall survival was evaluated using Kaplan-Meier method with survival distributions compared using log-rank test in 540 endometrial cancer patients from the Cancer Genome Atlas for HN1 stratified by tertile gene expression levels (log-rank p-value = 0.047).

**S1 - Supplemental Table 1:** Case identifiers, pre- and post-metformin treatment as well as metformin responder and non-responder status for endometrial cancer patient tissue samples analyzed.

**S2 - Supplemental Table 2:** Peptide spectral matches identified across endometrial cancer patient tissue samples analyzed.

**S3 - Supplemental Table 3:** Proteins significantly altered between responder versus non-responder patient tissue samples, pre-metformin treatment.

**S4A - Supplemental Table 4A:** Functional pathway analyses of proteins significantly altered between responder versus non-responder patient tissue samples, pre-metformin treatment – canonical pathways enriched.

**S4B - Supplemental Table 4B:** Functional pathway analyses of proteins significantly altered between Responder versus Non-responder patient tissue samples, pre-metformin treatment – functional pathways enriched.

**S5A - Supplemental Table 5A:** Proteins significantly altered between post versus pre-metfomrin treated tissues collected from metformin responders.

**S5B - Supplemental Table 5B:** Functional pathway analyses of proteins significantly altered between post versus pre-metformin treated tissues collected from metformin responders – functional pathways enriched.

**S6A - Supplemental Table 6A:** Proteins significantly altered between post versus pre-metfomrin treated tissues collected from metformin non-responders.

**S6B - Supplemental Table 6B:** Functional pathway analyses of proteins significantly altered between post versus pre-metformin treated tissues collected from metformin non-responders – functional pathways enriched.

**S7 - Supplemental Table 7:** HN1 IHC verification analyses.

**S8A - Supplemental Table 8A:** HN1 immunoprecipitation mass spectrometry analyses.

**S8B - Supplemental Table 8B:** Functional pathway analyses of HN1 immunoprecipitation mass spectrometry analyses – canonical pathways enriched.

